# Reference normalizations of Effort-To-Compress across length, alphabet size, and composition

**DOI:** 10.64898/2026.10.22.761192

**Authors:** Hyunwoo Jang, Jy-yong Sohn, Zirui Huang

## Abstract

Comparing finite symbolic complexity requires an explicit reference. Effort-To-Compress (ETC) counts the recursive pair substitutions needed to reduce a symbolic sequence to a constant. Its conventional normalization with *L* − 1 leaves systematic dependence on sequence length *L* and alphabet size *k*, confounding comparisons across datasets. We derive certified lower bounds for ETC and show that, for *L* ≥ *k* ≥ 2, the ceiling *L* − 1 is attainable if and only if *L* ≤ *k*^2^ + *k* + 1. For fixed *k* ≥ 2, the global ensemble mean and maximum ETC grow as Θ_*k*_(*L/* log_*k*_ *L*) as *L* → ∞. These support two normalized ETC coordinates. Global-ensemble normalized ETC (nETC) maps a certified lower bound to zero and the uniform ensemble mean at the same *L* and *k* to one. Composition-conditioned normalized ETC (cETC) uses corresponding references restricted to the observed symbol composition. In Potts chains, nETC substantially reduced dependence on length and alphabet size. cETC further reduced differences associated with unequal symbol frequencies, while retaining sensitivity to temporal dependence. In human chromosome 22, the exploratory predictor nETC – cETC outperformed LZ76 in distinguishing CpG islands and long interspersed nuclear elements from genomic background with AUROC of 0.947 and 0.848. Matched references therefore enable interpretable comparisons of symbolic complexity across sequence lengths, alphabet sizes, and compositions.

## I. INTRODUCTION

Comparing temporal structure of time series data is a recurring problem in statistical physics, signal processing, and neuroscience [1–3]. Compression-based measures quantify how readily repeated patterns reduce the resulting sequence [4, 5]. Among these methods, Effort-To-Compress (ETC) counts the Non-Sequential Recursive Pair Substitution (NSRPS) steps required to reduce a symbolic sequence to a constant [6, 7]. ETC has been applied to physiological recordings [8, 9] and human functional MRI [11, 12], where recording duration and quantization resolution can vary across acquisitions.

Raw ETC is a substitution count whose scale depends on both sequence length *L* and alphabet size *k*. It is conventionally divided by *L* − 1, the largest number of substitution steps allowed by sequence length alone [6, 11]. But obviously, this ceiling does not account for *k*, and the ceiling becomes increasingly unattainable as *L* grows (see Section III C). Consequently, ETC*/*(*L* − 1) drifts even when the same process is observed for a different number of samples or encoded at a different symbolic resolution (Fig. 1). Thus, the current convention cannot serve as a reliable common reference.

**FIG. 1.**
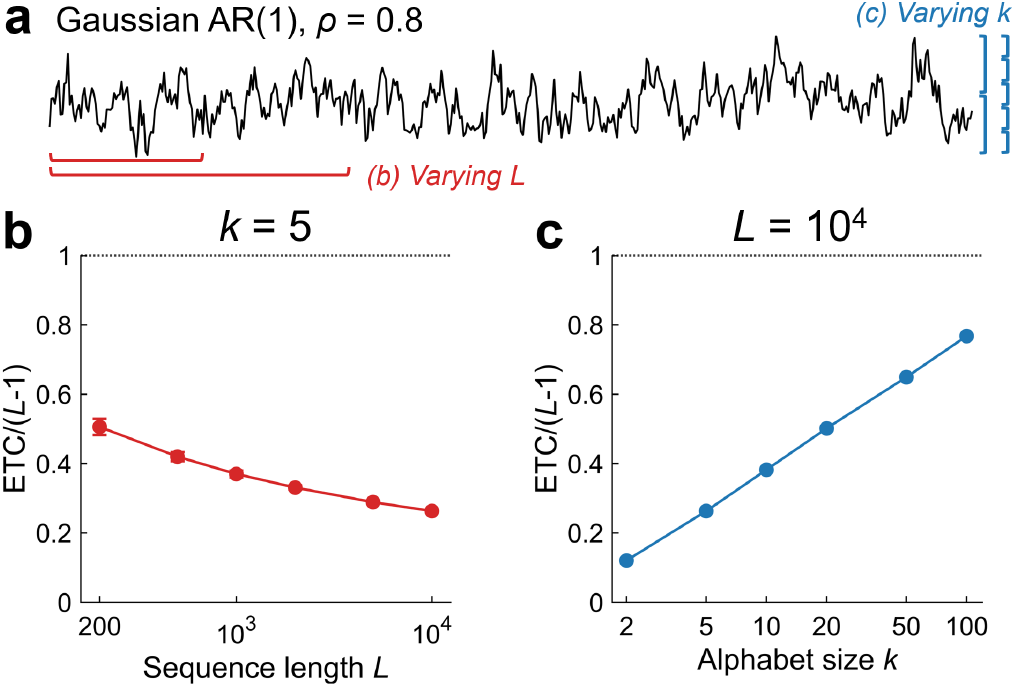
Dependence of conventionally normalized ETC on sequence length *L* and alphabet size *k*. (a) Representative stationary Gaussian first-order autoregressive [AR(1)] time series with *ρ* = 0.8. Red and blue brackets indicate the changes in *L* and *k* examined in (b) and (c), respectively. (b) ETC*/*(*L* − 1) decreases with *L* at fixed *k* = 5. (c) ETC*/*(*L* − 1) increases with *k* at fixed *L* = 10,000. Fifty independent time series of length 10,000 were generated and quantized to *k* equally populated bins. For each predefined *L* in (b), ETC was calculated from the first *L* samples of each time series. In (c), each full-length time series was re-quantized. Markers are mean *±* SD.

In this paper, we develop two normalized ETC coordinates. Following related reference-matching approaches for Lempel–Ziv (LZ) complexity [13, 14], global-ensemble normalized ETC (nETC) compares a given sequence with a certified lower bound and the uniform ensemble mean at the same *L* and *k*. Composition-conditioned normalized ETC (cETC) uses the corresponding references while matching the observed symbol counts and therefore controls the composition. We establish the lower bounds and their separation from the ensemble means to justify both normalizations. We also characterize when the conventional ceiling is attainable and how the ensemble mean and maximum ETC scale with sequence length. We use symbolic processes to evaluate whether nETC accounts for variation in *L* and *k*, and whether cETC additionally accounts for composition, while remaining sensitive to temporal structure. Finally, we apply nETC and cETC to annotated sequences from human chromosome 22 to examine what their different reference ensembles reveal about genomic organization.

## II. PRELIMINARIES AND REFERENCE ENSEMBLES

### A. Symbolic sequences and composition

For a symbolic sequence *S* = (*s*_1_, …, *s*_*L*_), let

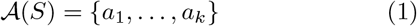

denote its observed alphabet, that is the set of distinct symbols appearing in *S*, where alphabet size *k* = |A(*S*)|. Unless stated otherwise, we assume *L* ≥ *k* ≥ 2. For each symbol *a*_*i*_, its multiplicity is

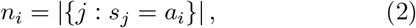

such that 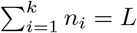. We call **n**_*S*_ = (*n*_1_, …, *n*_*k*_) the composition of *S*, and 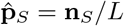 its empirical distribution. ETC(*S* ) is the number of recursive pair substitutions required to obtain a constant sequence. The formal algorithm and its implementation are given in Appendix A.

### B. Global and composition-conditioned ensembles

Let Ω_*L,k*_ denote the set of length-*L* sequences over a fixed ordered alphabet of *k* symbols, with every symbol appearing at least once. Its size is *k*!*S*(*L, k*), where *S*(*L, k*) is the Stirling partition number. We define ETC_min_(*L, k*), ETC_max_(*L, k*), and ETC_mean_(*L, k*) as the minimum, maximum, and uniform mean of ETC over the global ensemble Ω_*L,k*_.

For a positive labeled composition vector **n** = (*n*_1_, …, *n*_*k*_), let

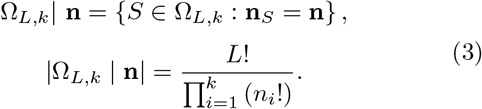

The notation | **n** restricts a reference quantity to sequences with composition **n**. For example, ETC_mean_(*L, k* | **n**) is the mean ETC over all distinct sequences that contain the prescribed number of each symbol.

Composition classes partition the global ensemble. Thus, unconditioned ETC_mean_(*L, k*) is the sequence-count-weighted average of their composition-conditioned means over all positive composition vectors.

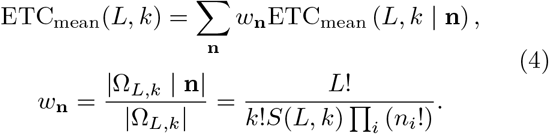

For example, at (*L, k*) = (4, 2), the global mean ETC is 2.714, while the composition classes (3, 1) and (2, 2) have mean ETC of 3 and 2.333, respectively. Thus, composition conditioning can increase or decrease the ensemble mean.

### C. Finite reference distributions

Fig. 2 compares global and composition-conditioned ETC distributions at *L* = (20, 1000) and *k* = 2. The global ensemble Ω_20,2_ contains 2^20^ − 2 = 1, 048, 574 sequences and was enumerated exhaustively. Its ETC distribution was 10.7 ± 1.1 (mean ± SD), ranging from 1 to 14 (Fig. 2(a,b)). The alternating sequences (01)^10^ and (10)^10^ attained the minimum. Forty-eight distinct sequences attained the maximum, including 10011110101000000101. Conditioning on **n** = (15, 5) selected ^20^*C* _5_ = 15,504 sequences and shifted the distribution to 10.0 ± 1.0, ranging from 3 to 13.

**FIG. 2.**
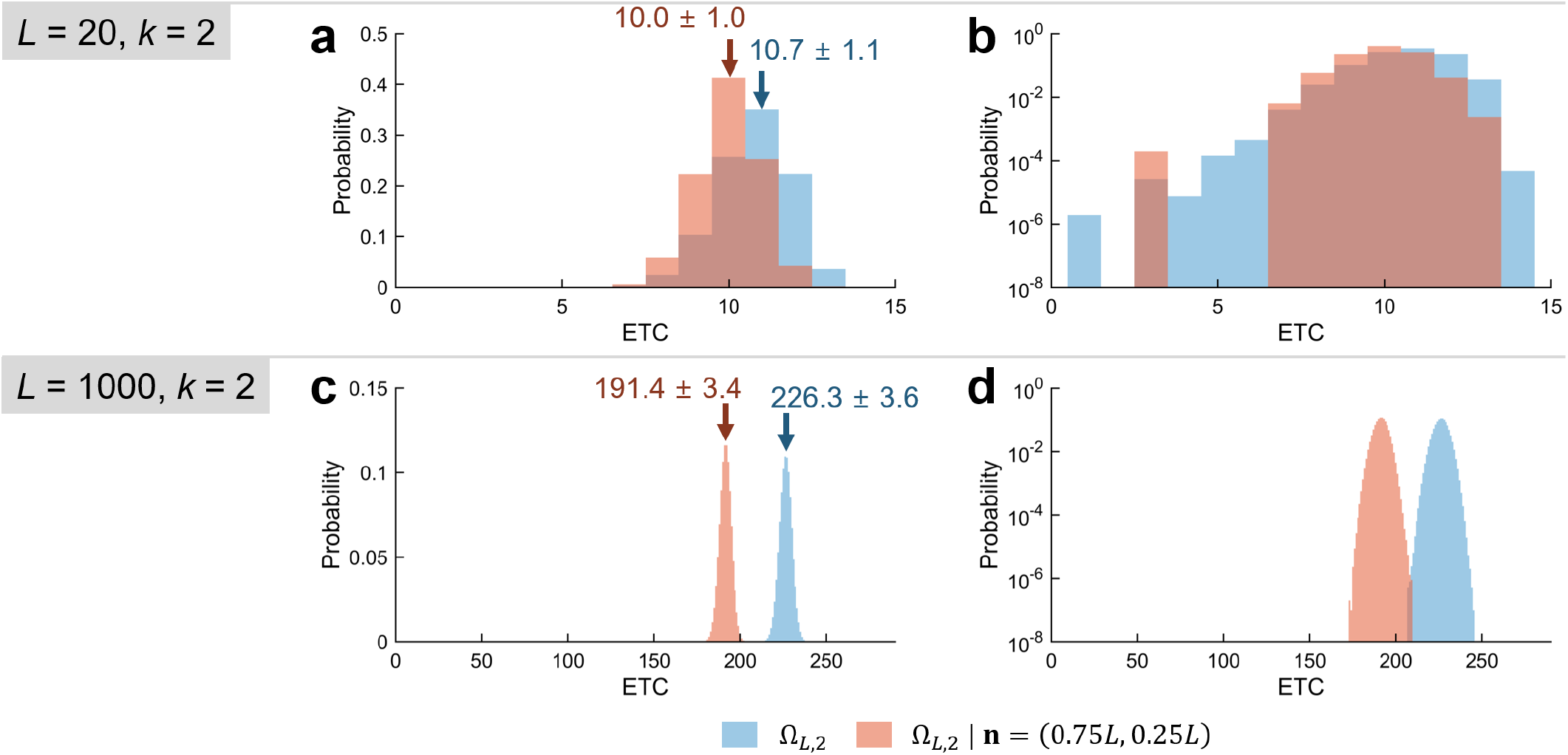
Global and composition-controlled ETC distributions. (a,b) Exact probability distribution of ETC over Ω_20,2_ and its composition-controlled subset Ω_20,2_ | **n** = (15, 5). (c,d) Empirical probability distributions of Ω_1000, 2_ and Ω_1000,2_ | **n** = (750, 250) from 10^7^ uniform Monte Carlo samples. Blue and orange denote the global and composition-conditioned ensembles, respectively. Panels a,c are in linear scale, while b,d are in logarithmic scale. Numbers in panels a,c are mean *±* SD.

Ω_1000, 2_ contains 2^1000^ − 2 sequences which cannot be fully enumerated. We therefore estimated the global and composition-conditioned distributions using 10^7^ uniform samples from each ensemble. The estimated mean was at 226.3 ± 3.6 for the global ensemble and 191.4 ± 3.4 under **n** = (750, 250) (Fig. 2(c,d)). Their sampled ranges were 207–245 and 173–209, respectively. Because each composition-conditioned ensemble is a subset of the corresponding global ensemble, conditioning can only preserve or narrow the theoretical ETC range. Together, the exact and sampled distributions show that composition can change the reference mean and range. Ensemble sampling rules used in this paper are given in Appendix A.4.

## III. NORMALIZED ETC AND REFERENCE ANCHORS

### A. Two normalized ETC coordinates

The global and composition-conditioned ensembles define two normalized coordinates of ETC. Global-ensemble normalized ETC is

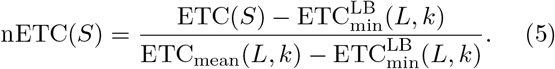

where 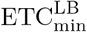denotes a certified lower bound satisfying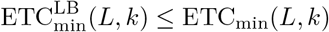, since exact minimum ETC is not generally available (see Section III.B).

Therefore, nETC maps the lower bound to zero and the global ensemble mean to one at fixed (*L, k*). Because global ensemble includes all possible compositions, differences in symbol multiplicity remain part of the measured outcome.

Composition-conditioned normalized ETC is

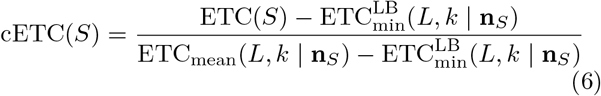

cETC compares the observed sequence *S* with the subset of global ensemble conditioned to the composition of empirical sequence *S*.

Thus, nETC measures composition and symbol ordering relative to the global ensemble, whereas cETC only evaluates ordering structure while controlling symbol multiplicity. nETC and cETC can therefore answer different comparison questions.

### B. Certified lower bounds and well-definedness of ensemble normalization

The choice of lower anchor determines the calibrated range of analytically justified ETC. A valid lower anchor must not exceed the ETC of any sequence in the corresponding reference ensemble. Zero is a universal bound, but it is unattainable within Ω_*L,k*_ for *k* ≥ 2. A sharper trivial bound is *k* − 1, because one substitution can reduce the number of active symbols by at most one. When *k* divides *L*, this bound is attained by the periodic sequence (0, 1, …, *k* − 1)^*L/k*^ . We derive tighter closed-form anchors by incorporating the constraints imposed by *L* and composition **n**_*S*_.

Let *M* (*L, k*) be the largest divisor of *L* that does not exceed ⌊*L/k*⌋. For a positive composition vector **n**_*S*_ = (*n*_1_, …, *n*_*k*_), define *g*(**n**_*S*_) = gcd(*n*_1_, …, *n*_*k*_).

#### Proposition 1

**(Certified lower bounds on ETC)** *For integers L* ≥ *k* ≥ 2, *every sequence S* ∈ Ω_*L,k*_ *satisfies the global bound*

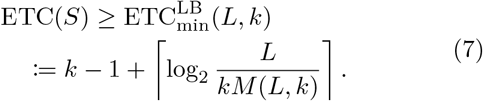

*For its composition vector* **n**_*S*_, *the composition-conditioned bound is*

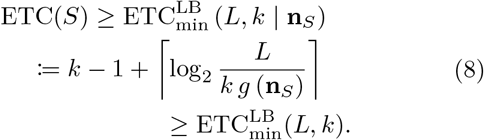

*These bounds obey*

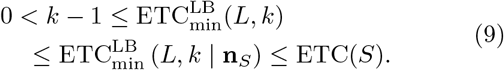

The reverse-substitution proofs of Proposition 1 are given in Appendices B.1 and B.2. The global anchor depends only on *L* and *k*. When *k* divides *L, M* (*L, k*) = *L/k*, so the anchor reduces to *k* − 1. Conditioning on composition can further tighten the anchor.

The normalized coordinates are well defined only if each random-mean anchor lies strictly above its corresponding lower anchor.

#### Proposition 2

**(Strict anchor separation** *For integers L* ≥ 4 *and* 2 ≤ *k* ≤ *L* − 2, *the global ensemble mean satisfies*

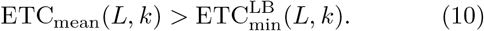

*For every positive integer composition vector* **n**_*S*_ = (*n*_1_, …, *n*_*k*_) *with*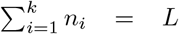, *the composition-conditioned mean satisfies*

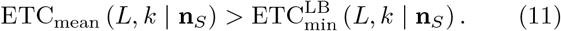

Appendix B.3 proves Proposition 2. Thus, the denominators of nETC and cETC are positive throughout the stated domain. At *k* = *L*−1 or *k* = *L*, both denominators vanish. The case *k* = 1 contains only constant sequences with ETC zero.

For nETC, replacing the true minimum with the certified lower bound changes the normalized value by at most *O*_*k*_ ((log^3^ *L*)*/L* at fixed *k*, as shown in Proposition 4 (Appendix B.4).

### C. Upper scale and fixed-*k* asymptotics

Values of nETC above one indicate substitution counts above the global ensemble mean. Such values are possible because the mean is a reference, not a ceiling. The conventional normalization denominator *L* − 1 is a trivial ceiling for ETC, but it is not always attainable.

#### Proposition 3

**(Exact ceiling threshold)** *For integers L* ≥ *k* ≥ 2, *let T* (*k*) = *k*^2^ + *k* + 1. *Then*

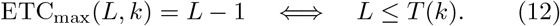

Appendix C proves Proposition 3 and constructs a sequence attaining *L* − 1 for every *k* ≤ *L* ≤ *T* (*k*). For *L > T* (*k*), even the maximum ETC is strictly below *L* − 1.

#### Theorem 1

**(Fixed-*k* ETC scaling)** *For each fixed integer k* ≥ 2, *the global-ensemble mean ETC and maximum ETC satisfy, as L* → ∞,

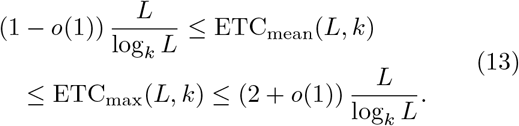

*In particular, both quantities have order* Θ_*k*_ (*L/* log_*k*_ *L*).

Appendices D.1 and D.2 prove the lower and upper bounds in Theorem 1, respectively. These bounds do not determine exact leading coefficients.

Theorem 1 shows that the ensemble mean and maximum ETC grow at the same sublinear order for fixed *k*, while the conventional denominator remains linear in *L*.

#### Corollary 1

**Conventional normalization)** *For each fixed integer k* ≥ 2, *the mean conventionally normalized ETC satisfies as L* → ∞

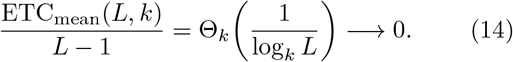

Dividing the bounds in Theorem 1 by *L*−1 proves Corollary 1 because *L/*(*L* − 1) → 1. Thus, the conventional normalization sends the global reference mean toward zero as sequence length increases.

Theorem 1 also bounds the maximum-to-mean ratio:

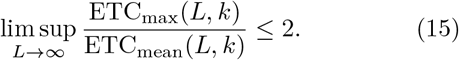

This is a bound on the limiting upper ratio. Whether the ratio converges or the global mean asymptotically approaches the maximum is unknown.

The same order applies to composition-conditioned means. Fix an integer *k* ≥ 2 and a positive symbol distribution **p** = (*p*_1_, …, *p*_*k*_) that sums to one. Let **n**^(*L*)^ be a positive integer composition with 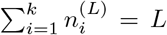 and **n**^(*L*)^ */L* → **p**. Define

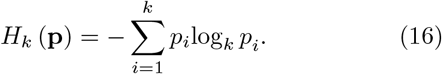

Then

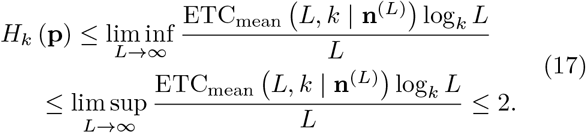

Proposition 5 (Appendix D.3) establishes these bounds. The lower bound follows by counting the binary grammars that can represent sequences with a given ETC. The upper bound holds for every sequence and therefore also applies to each composition-conditioned mean.

### D. Sequences above the global ensemble mean

We constructed sequences that achieve nETC > 1 using a greedy search. The search starts from the de Bruijn-based threshold construction in Appendix C. This sequence has length *L* = *T* (*k*) and ETC = *T* (*k*) − 1. The search then tests symbol insertions in a fixed order and retains a candidate when the newly computed ETC increases. If no one-symbol insertion increases ETC, the search tests short-block insertions. If these also fail, it retains the one-symbol candidate with the highest evaluated ETC. Although the search is heuristic, the ETC of each constructed sequence provides a certified lower bound on the maximum ETC. Therefore, for each greedy-searched sequence *G*_*L,k*_,

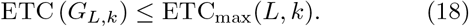

Appendix E gives the full algorithm.

Fig. 3 shows representative ETC trajectories of *G*_*L,k*_ for *k* = 2, 5, 10, 20, and 50. We estimated the global ensemble mean from 200 uniform Monte Carlo samples at each selected (*L, k*). After division by *L* − 1, both the greedy ETC values and the estimated ensemble means decreased overall with *L* (Fig. 3(a)). At the sampled reference lengths, each greedy ETC value exceeded its corresponding estimated ensemble mean. On the global-ensemble scale, this separation appeared as nETC > 1 (Fig. 3(b)). For sampled lengths *L* > 100, nETC (*G*_*L,k*_) ranged from 1.12 to 1.34 with visible undulations. The asymptotic limit of nETC (*G*_*L,k*_) remains unknown.

**FIG. 3.**
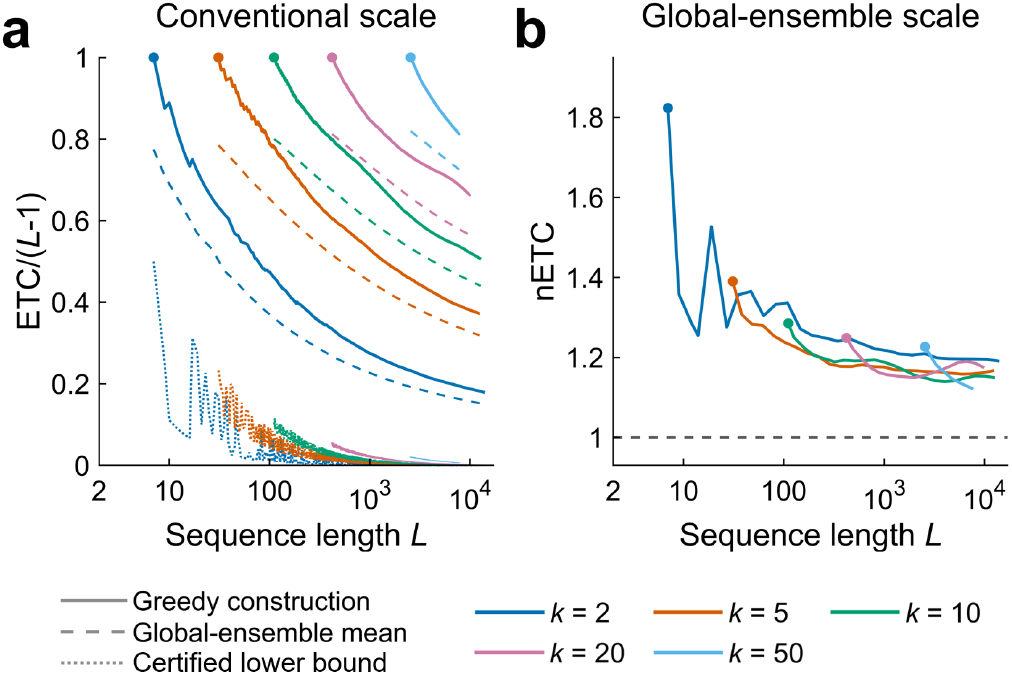
Greedy-constructed sequences above the estimated global ensemble mean. (a) Solid, dashed, and dotted curves show the ETC of greedy-constructed sequences, the estimated global ensemble mean, and the certified global lower bound, respectively, each divided by *L* − 1. Curves begin at *T* (*k*) = *k*^2^ + *k* + 1. Ensemble means were estimated from 200 uniform Monte Carlo samples per (*L, k*). (b) nETC of the constructed sequences at the sampled reference lengths. Lines connecting Monte Carlo-based values serve as visual guides. The horizontal line marks nETC = 1. Filled circles mark the exact linear-ceiling thresholds *T* (*k*).

## IV. CONTROLLED CALIBRATION IN POTTS CHAINS

We evaluated nETC and cETC using a one-dimensional *k*-state Potts chain represented as a stationary finite-state Markov process [15, 16]. Two parameters, *J* and *η*, were used as generative parameters. The nearest-neighbor coupling *J* primarily modulates one-step temporal dependence, while one-symbol field *η* introduces compositional bias to single favored symbol. When *J* = 0, the symbols are independent and identically distributed (iid) for any *η*.

We parameterized temporal dependence using the normalized entropy rate 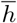 defined as

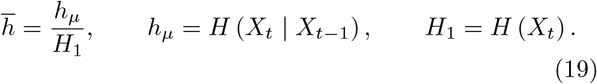

where *X*_*t*_ denotes the symbol at position *t*. The numerator *h*_*µ*_ is the entropy rate of the stationary first-order Markov process, and *H*_1_ is its symbol entropy. When all symbols are equiprobable, i.e., in uniform composition, *H*_1_ = ln *k*. Under an iid condition, *h*_*µ*_ = *H*_1_ and therefore 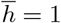. Temporal dependence reduces *h*_*µ*_ relative to *H*_1_, giving 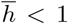. Thus, 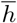 measures transition uncertainty relative to the uncertainty of the corresponding marginal composition. This definition allows the same scale of temporal dependence under uniform and nonuniform compositions.

Here, uniform composition refers to uniform stationary symbol probabilities, not equal empirical counts in every finite sequence.

For the nETC benchmark, zero-field chains (*η* = 0) were used across sequence lengths and alphabet sizes under uniform composition. For the cETC calibration, we specified favored-symbol probability *p*_*f*_ and normalized entropy rate 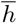, then solved jointly to get corresponding *J* and *η*. See Appendix F for detailed definitions of the Potts parameters.

### A. Length and alphabet-size calibration of nETC

We first examined the uniform-composition case. We compared conventional ETC*/*(*L* − 1) and nETC across sequence lengths *L* = 2,000–16,000 and alphabet sizes *k* = 2–50 (Fig. 4(a)).

**FIG. 4.**
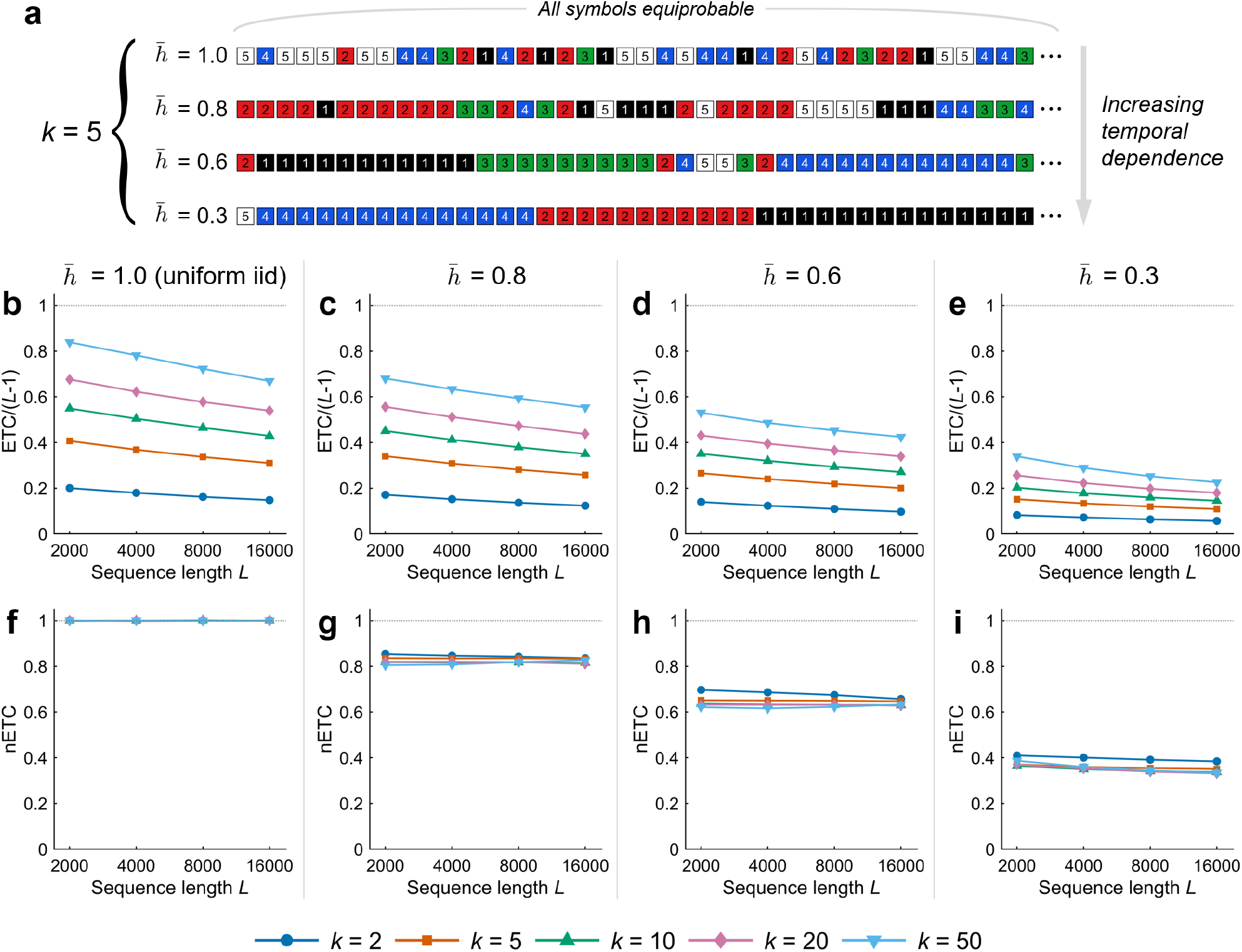
Joint calibration of nETC across sequence length and alphabet size. (a) Representative 40-symbol excerpts from zero-field Potts sequences of varying normalized entropy rates 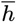 at fixed *k* = 5. 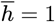 corresponds to uniform iid condition and decrease in 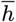 increases one-step temporal dependence. (b–e) Conventionally normalized ETC and (f–i) nETC at 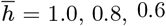, and 0.3. Curves show means over 200 sequences. The nETC reference means were also estimated from 200 sequences drawn uniformly from the global ensemble at the same *L* and *k*. Horizontal dotted lines indicate the conventional ceiling in (b–e) and the global-ensemble mean in (f–i).

Conventional ETC*/*(*L* − 1) depended strongly on both *L* and *k*, varying from 0.15 to 0.84 across the *L–k* grid at 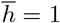 (Fig. 4(b)). It decreased with larger *L* and smaller *k*. The same trends persisted at lower entropy rates (Fig. 4(c– e)). By contrast, nETC showed substantially less variation with *L* and *k* while retaining the sensitivity to temporal dependence (Fig. 4(f–i)). Under iid condition 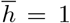 Fig. 4(f)), nETC remained approximately one throughout the grid. At 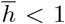, nETC consistently decreased with 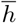(Fig. 4(g–i)), whose values closely followed their corresponding normalized entropy rates except for 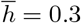. Their proximity was not imposed by normalization, and the asymptotic relationship is unknown.

### B. Composition calibration of cETC

We next examined whether cETC controls symbol composition while retaining sensitivity to temporal dependence. We generated five-state Potts sequences of length *L* = 2,000 at favored-symbol probability *p*_*f*_ = 0.2, 0.35, 0.5, 0.65, and 0.8 and normalized entropy rate 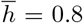, 0.85, 0.9, 0.95, and 1.0 (Fig. 5(a–d)).

**FIG. 5.**
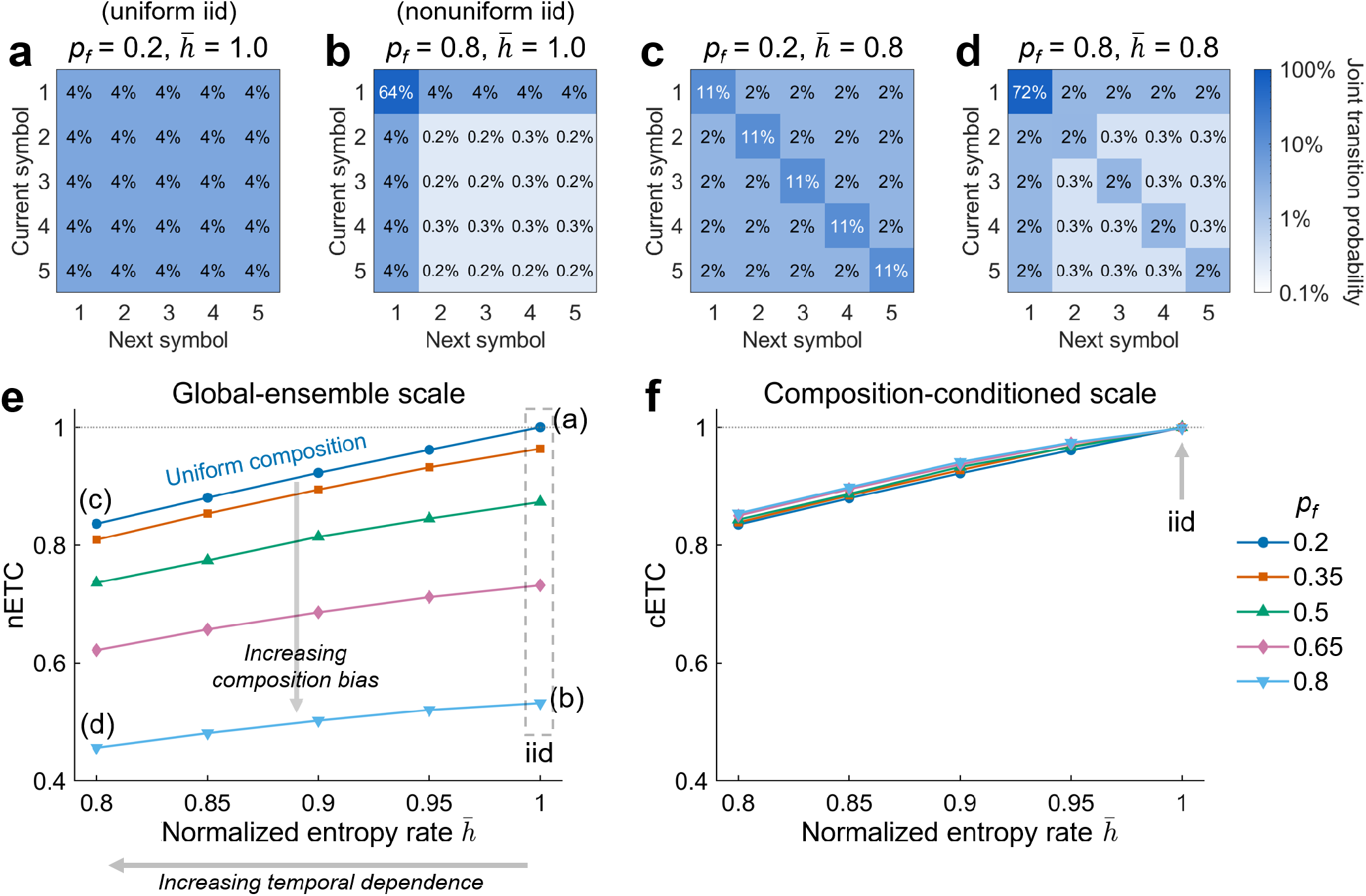
Composition calibration of cETC in five-state Potts chains. (a–d) Joint transition probabilities for four representative combinations of favored-symbol probability *p*_f_ and normalized entropy rate *h* = *h*_*µ*_*/H*_1_ . Here, symbol 1 is favored. Panels (a) and (b) show uniform and nonuniform iid processes; panels (c) and (d) add one-step temporal dependence at the respective compositions. (e,f) nETC and cETC across *p*_*f*_ = 0.2–0.8 and *h* = 0.8–1.0 for sequences of length *L* = 2,000. Corresponding points for panels (a–d) are denoted. *p*_*f*_ = 0.2 represents uniform composition for *k* = 5. Higher *p*_*f*_ and smaller *h* correspond to increase in compositional bias and temporal dependence, respectively. Curves show means over 200 sequences. Reference means were estimated from 200 samples.

Under iid conditions 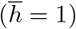, nETC decreased from 1.0 to 0.53 as *p*_*f*_ increased from 0.2 to 0.8 (dashed rectangle in Fig. 5(e)). Thus, unequal symbol occurrence dropped nETC even with no temporal dependence. Over the same compositions, the cETC remained approximately 1.0 (gray arrow in Fig. 5(f)). This shows that composition conditioning aligned these iid processes with their respective ensemble references.

As 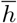 decreased and temporal dependence increased, both nETC and cETC decreased monotonically while the curves for different compositions remained close for cETC (Fig. 5(e,f)). Composition matching therefore reduced differences associated with symbol distribution while preserving the response to temporal dependence.

## V. APPLICATION TO GENOMIC SEQUENCE CLASSIFICATION

We applied nETC and cETC to human chromosome 22 sequences of Genome Reference Consortium Human Build 38 (GRCh38) and examined whether the two reference ensembles help distinguish genomic annotations from background. Normalized LZ76 (nLZC) was included for comparison based on its previous applications in genomic analysis [10, 17]. We selected CpG island, long interspersed nuclear element (LINE), and coding exon as target annotations (Fig. 6(a)). Each sequence was paired with a background sequence of the same length. We evaluated three separate binary classification tasks, each comparing one annotation with its matched background. See Appendix G for methodological details.

**FIG. 6.**
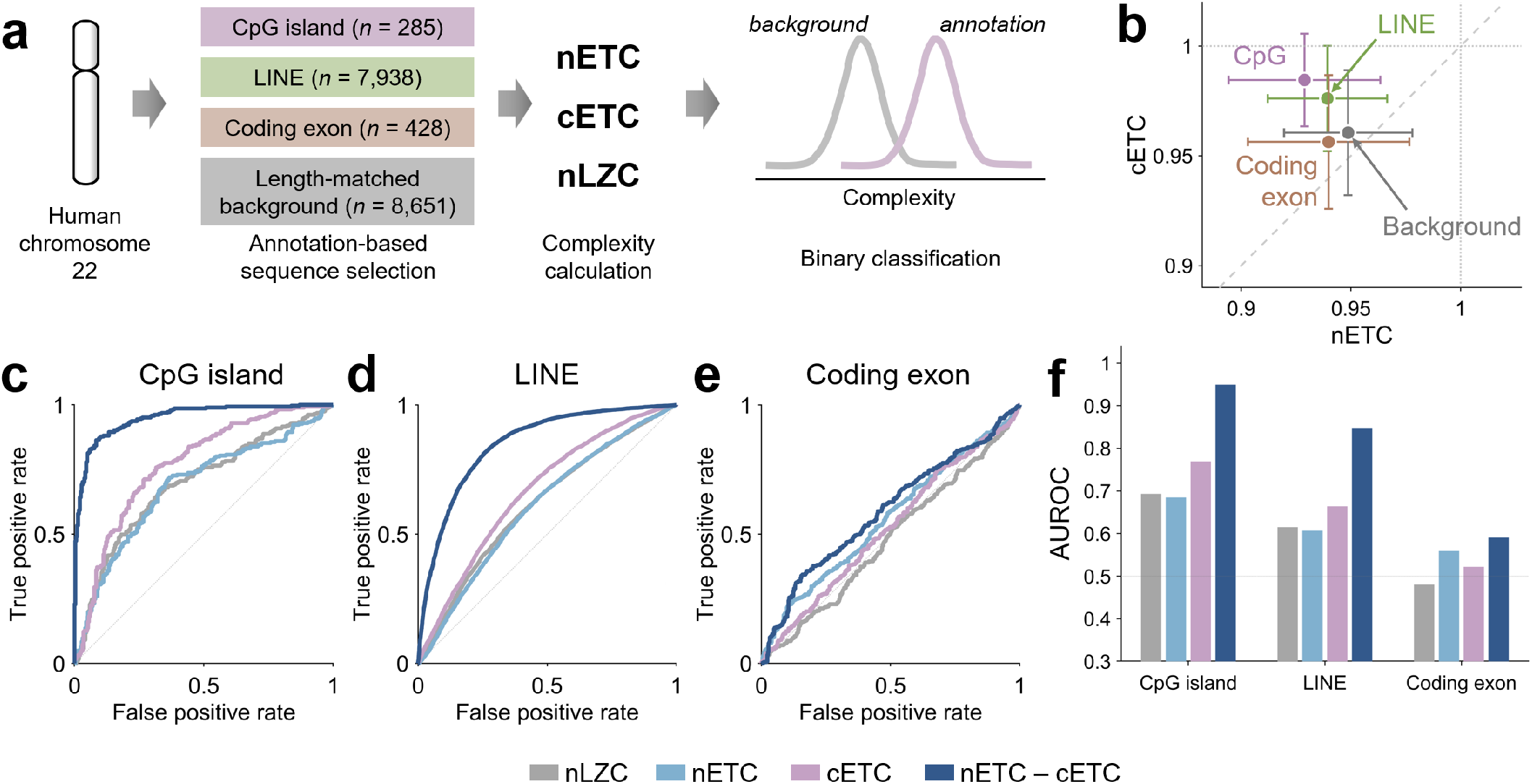
Complexity-based discrimination of genomic annotations in human chromosome 22. (a) Analysis workflow for CpG island, long interspersed nuclear element (LINE), coding exon, and length-matched background sequences from human chromosome 22. Each annotation was classified against its matched background. (b) Mean nETC and cETC with horizontal and vertical SD bars. For visualization, background sequences are pooled across the three annotation groups. The diagonal indicates nETC = cETC. (c–e) Receiver operating characteristic (ROC) curves for CpG islands, LINEs, and coding exons using normalized LZ76 (nLZC), nETC, cETC, or nETC *−* cETC as a single predictor. Curves pool out-of-fold predictions from logistic regression under five-fold grouped cross-validation. (f) Corresponding areas under the ROC curves (AUROCs).

After adjustment for sequence length, CpG islands and LINEs had lower nETC (*β* = −0.0207 and −0.00949, respectively; *p*_FDR_ < 0.05; two-sided *t* tests of the annotation coefficients) but higher cETC (*β* = 0.0246 and 0.0155, respectively; *p*_FDR_ < 0.05) than their matched backgrounds (Fig. 6(b)). nLZC was also lower in both annotations (*p*_FDR_ < 0.05). See Table G1 for full statistics. We evaluated each feature as a single predictor in logistic regression with five-fold grouped cross-validation. The opposite shifts in nETC and cETC motivated the exploratory inclusion of nETC − cETC as a single predictor. The difference nETC − cETC yielded high AUROCs of 0.947 and 0.848 for CpG islands and LINEs, respectively (Fig. 6(c,d,f)). Other complexity metrics yielded AUROCs of 0.61–0.77 for CpG islands and LINEs. Thus, nETC − cETC provided greater discrimination of CpG islands and LINEs from their matched backgrounds than either coordinate alone or nLZC.

Coding exons had lower nETC and nETC − cETC than their matched background (*β* = −0.00767 and −0.00524; *p*_FDR_ < 0.05). The AUROC achieved by nETC − cETC was 0.597 (Fig. 6(e,f)). A detectable mean difference therefore did not provide strong discrimination of individual coding exons. Neither cETC nor nLZC showed a significant difference (*p*_FDR_ > 0.05; Fig. 6(b)).

## VI. DISCUSSION

Normalization choice is a recurring problem in compression-based complexity. In LZ complexity, conventional normalization uses the asymptotic scale *L/* log_*k*_ *L* while alternative approaches such as an exact finite-length maximum [18] and normalization against shuffled sequences were proposed [19]. Our framework of an ensemble-based normalization of ETC is similar to the latter approach, but also combines certified lower bounds. By accounting for length and alphabet size, ensemble-based normalization enables more consistent comparisons of temporal structure, as demonstrated in the Potts experiments.

Composition conditioning addressed another important ambiguity in interpreting ETC complexity. A sequence dominated by one symbol contains frequent repetitions even when the ordering of symbols is random and has no temporal dependence. cETC compares the observed sequence with random rearrangements with exactly the same symbol counts and asks whether the observed ordering requires fewer substitutions. This follows the logic of constrained surrogate data, which preserve selected properties while randomizing others [20]. cETC therefore distinguishes repetition expected from symbol abundance from additional compressibility associated with their ordering structure.

Jointly interpreting nETC and cETC can reveal distinctions that either coordinate alone misses. In the exploratory classification analysis, the heuristic contrast nETC − cETC yielded higher AUROC for CpG island and LINE than either coordinate alone or normalized LZ76. This contrast describes how conditioning on composition changes the assessment of compression resistance. A negative contrast indicates that the sequence appears more compressible against the global reference than against its composition-matched reference. When cETC remains near one, a low nETC is therefore consistent with a composition-associated reduction rather than unusually compressible ordering. A positive contrast indicates that composition matching makes the sequence’s compressibility more pronounced relative to its reference. A negligible contrast indicates agreement between the two normalized assessments while it does not necessarily establish uniform composition. Because both the reference means and lower anchors can differ, the contrast does not quantify the proportion of complexity attributable to composition. Further work could establish when this subtraction has mathematical foundation and clarify how its behavior relates to symbolic properties.

Randomness and resistance to a compression algorithm are distinct properties of finite sequences. A deliberately arranged sequence can resist compression by limiting the repetitions that an NSRPS substitution can remove. High ETC complexity thus need not arise from a random generation rule but can also arise from an arrangement designed to accomplish little compression. Prior work showed that deterministic constructions can attain higher finite LZ complexity than typical random sequences [18]. Our greedy construction pursued such compression resistance by favoring insertions with smaller maximum adjacent-pair counts (Figs. 3 and E1). By using ensemble mean as an upper reference, the proposed ETC scales can directly reveal the supra-random compression resistance of a sequence (nETC, cETC > 1).

The connection between NSRPS and entropy rate has been discussed [21, 22]. Intriguingly, our Potts experiments show that ETC, as a stopping-count measure, also tracks normalized entropy rate after reference normalization. The numerical agreement across lengths and alphabet sizes supports the proposed calibration, particularly because entropy rate never enters the definitions of nETC or cETC. It may also reflect a deeper relationship between substitution count and entropy rate that existing NSRPS theory does not establish.

This possibility raises a further question: do nETC and cETC primarily reflect unpredictability, or can they also distinguish processes with the same entropy rate but different memory requirements? In the first-order Potts chains, the previous symbol contains all the history needed for optimal prediction. But processes with the same entropy rate can require different amounts of predictive history [23]. Recursive pair substitution can build symbols representing progressively longer patterns. Whether the final substitution count captures differences in predictive memory remains to be tested. Comparing processes with matched entropy rates, lengths, and marginal symbol probabilities but different memory requirements would help determine what normalized ETC captures beyond entropy rate.

Our work has limitations. Exact closed-form formula or computationally tractable algorithm to obtain minimum, mean, and maximum ETC for any *L* and *k* remain unresolved, warranting future investigation. The applicability to continuous time series recordings is to be validated. In such cases, measurement choices including sampling interval and quantization methods (e.g., the choice of *k* and bin width selection) can significantly impact symbol composition and temporal ordering. Whether nETC and cETC are immune to these factors is unknown. Future work should investigate these effects to guide comparisons across time series data.

In conclusion, matched lower and ensemble-mean references provide interpretable ETC scales across sequence lengths, alphabet sizes, and observed symbol counts. Together, these coordinates provide complementary views of finite symbolic structure.

## ACKNOWLEDGMENTS

This work was supported by the National Institute of General Medical Sciences of the National Institutes of Health (grant R01GM103894; PIs: Anthony G. Hudetz and Zirui Huang) and the Center for Consciousness Science, Department of Anesthesiology, University of Michigan Medical School, Ann Arbor, Michigan, USA. The content is solely the responsibility of the authors and does not necessarily represent the official views of the National Institutes of Health.

## DATA AVAILABILITY

The source data are available from the corresponding author upon request. The C implementation of ETC is publicly available at https://github.com/janghw4/normalized-effort-to-compress. The genomic reference sequence and annotations are available from the sources cited in Appendix G.

## Appendix A: ETC algorithm and its implementation

### 1. Substitution rule and stopping convention of ETC

ETC counts the pair-substitution steps required to reach a constant sequence [6]. We follow the pair-selection and substitution conventions of the authors’ MATLAB implementation [24], with exact constant-sequence termination.

Let *S* = (*x*_1_, …, *x*_↕_) be the current sequence. Symbols have a fixed numerical order. For an ordered pair (*a, b*), let *f*_*S*_(*a, b*) be the number of nonoverlapping occurrences selected by a left-to-right scan. After a match at positions *i, i* + 1, the scan resumes at *i* + 2. Otherwise it resumes at *i* + 1. Define

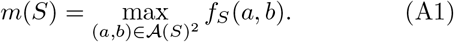

If *S* is not constant, select a pair with frequency *m*(*S*). Among tied pairs, select the smallest second label and then the smallest first label. Replace all selected occurrences by one fresh symbol whose label exceeds every existing label. Repeat until the sequence is constant. The number of substitutions is ETC(*S*). A nonempty constant input has ETC zero. The order of the original labels is part of this deterministic convention.

Each substitution defines a binary grammar rule and reduces the sequence length by its selected nonoverlapping frequency. The terminal constant contributes no additional steps. Therefore,

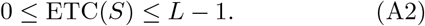

### 2. Frequency-one regime

If the maximum nonoverlapping pair frequency becomes one before termination, the sequence enters the frequency-one regime.

#### Lemma 1

**(Frequency-one regime)** *Let S be a non-constant current sequence of length p with m*(*S*) = 1. *The maximum nonoverlapping pair frequency remains one at every subsequent nonconstant step, and exactly p* − 1 *further substitutions are required to reach a constant sequence*.

Each substitution replaces a single occurrence and creates a fresh symbol that occurs once. Every newly formed pair contains that symbol and is therefore unique, while no existing pair count increases. Each subsequent substitution thus reduces the length by exactly one. The newly created symbol also prevents the sequence from becoming constant while its length exceeds one. Termination therefore occurs at length one, after exactly *p* − 1 further substitutions.

Not every sequence enters this regime. For example, 0101 becomes the constant sequence 22 after one substitution of a pair with frequency two. The shortcut applies only when the current sequence is nonconstant and its maximum pair frequency is one. Once this condition holds, the remaining *p* – 1 steps can be counted without performing the substitutions.

### 3. C implementation

The C implementation uses integer symbol labels and an array of pair counts. It preserves the label order and the tie-breaking rule of original implementation [24]. Each substitution scans the sequence from left to right and writes the result to a second buffer. The two buffers exchange roles after each step. The implementation checks whether the sequence is constant before counting pairs. It therefore does not calculate Shannon entropy or use a numerical entropy threshold to determine termination. When the maximum pair frequency is one, it adds the current length minus one to the substitution count and returns, using Lemma 1. On a benchmark corpus of 16,607 sequences, the C implementation returned identical ETC values to the reference MATLAB implementation. Median execution time across three runs was 0.038 s for C and 2.690 s for MATLAB on an AMD Ryzen 9 9950X3D processor, corresponding to a 71-fold speedup.

### 4. Ensemble sampling rules

The global ensemble Ω_*L,k*_ was defined on a fixed ordered alphabet of *k* symbols and contains only sequences in which every symbol occurs. A global reference can be sampled exactly by drawing iid uniform symbols and rejecting sequences that omit any alphabet symbol. The conditioning is required even when rejection is rare.

Global references were generated using either rejection sampling or the completion-count sampler described below. Let *D*(*r, u*) be the number of valid sequences when *u* unseen symbols are remaining after filling in *L* − *r* positions. Then

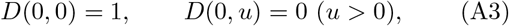

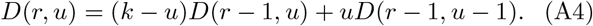

Choose the unseen-symbol category with probability *uD*(*r* − 1, *u* − 1)*/D*(*r, u*) and otherwise choose the seen-symbol category. Within the chosen category, select a symbol uniformly. This guarantees equal probability to every member of Ω_*L,k*_.

The composition-conditioned ensemble was estimated by counting the symbol occurrences and constructing a reference sequence containing exactly *n*_*i*_ copies of symbol *a*_*i*_. We then randomly shuffled all *L* positions and computed ETC. Repeating this procedure estimated the composition-conditioned mean. Original numerical labels were not sorted or reassigned.

## Appendix B: Certified lower references and anchor separation

### 1. Global-ensemble certified lower bound

Suppose ETC(*S*) = *e* and the terminal constant (i.e., the constant sequence after the last NSRPS step) has multiplicity *v*. Reverse *e* substitutions from the terminal word *a*^*v*^. We call the resulting sequence *P* the expansion of *a*. Every copy of *a* expands through the same grammar, so *S* = *P*^*v*^ for one word *P* . Hence *v* | *L*. Every original symbol occurs a positive multiple of *v*, which also gives *v* ≤ ⌊*L/k*⌋.

Let *Q*_*i*_ be the largest symbol multiplicity after completing *i* reverse expansion steps. Each expansion step changes the number of current symbols by +1, 0, or −1.

- +1: Two distinct new symbols are created. *Q*_*i*_ does not increase.
- 0: *Q*_*i*_ increases by at most a factor of two.
- −1: Two parental multiplicities can be added to an existing child. Thus *Q*_*i*+1_ ≤ 3*Q*_*i*_ ≤ 4*Q*_*i*_.

Let *p, z*, and *q* count these three step types among *e* expansion steps (*e* = *p* + *z* + *q*). Since the symbol count grows from one to *k* during the entire expansion, *p* − *q* = *k* − 1. Thus

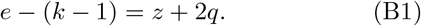

Starting from *Q*_0_ = *v* gives

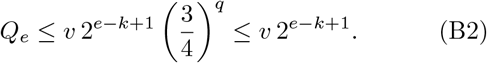

Since *Q*_*e*_ is the largest symbol multiplicity of the original sequence *S, L* ≤ *kQ*_*e*_.

Let *M* (*L, k*) be the largest divisor of *L* not exceeding ⌊*L/k*⌋. Since *v* ≤ *M* (*L, k*), taking the integer ceiling gives

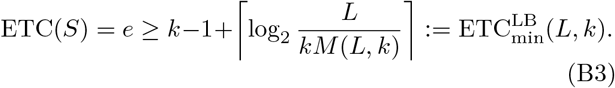

### 2. Composition-conditioned certified lower bound

For a fixed positive composition **n**_*S*_, the identity *S* = *P*^*v*^ implies that *v* divides every *n*_*i*_. Therefore, *v* ≤ *g* (**n**_*S*_) = gcd (*n*_1_, …, *n*_*k*_). Applying the multiplicity bound from Appendix B.1 gives

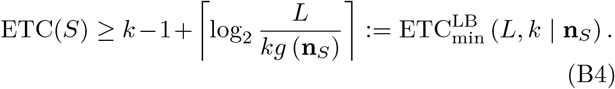

### 3. Strict separation of ensemble means and lower bounds

Here, we prove that there is at least one sequence that guarantees larger ETC than composition-conditioned lower bound for any admissible **n**_*S*_ for 2 ≤ *k* ≤ *L* − 2. Every other sequence has ETC at or above that bound. The ensemble mean must therefore lie strictly above it. The main task is to construct such a sequence for each composition.

#### a. Counting substitutions in a grouped-run sequence

First place all copies of each symbol in grouped-run structure:

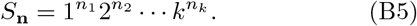

This sequence matches the required composition **n**. Pairs that cross a boundary of symbol blocks (e.g., 12, 23, … ) occur only once. As long as any pair occurs at least twice, the algorithm selects a pair inside a block. Substitutions in one block do not change the contents of the other blocks.

Within each symbol block, a substitution halves the leading run and leaves one unpaired symbol if its length is odd. For example, a block of 13 copies of A changes as *A*^13^ → *B*^6^*A* → *C*^3^*A*, where *B* and *C* are fresh symbols. After these two substitutions, four symbols remain where the nonoverlapping frequency of every pair is one.

More generally, for a block of length *x* ≥ 2, let *t* = ⌊log_2_ *x*⌋. Halving the leading run until its length is two or three takes *t* − 1 substitutions. Each odd-length halving leaves one symbol behind. These remainders correspond to the lower binary digits of *x*, while the final run contains two or three symbols. The residual length is therefore wt_2_(*x*) + 1, where wt_2_(*x*) is the Hamming weight of *x*, i.e., number of ones in its binary expansion. Define

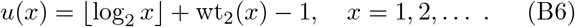

Thus, *u*(*x*) equals the number of these substitutions plus the residual length minus one. In the example of *A*^13^, *u*(13) = 2 + 4 − 1 = 5, which is also two substitutions plus four residual symbols minus one. This quantity describes a block within the nonconstant grouped-run sequence.

Let *h*_*i*_ be the number of substitutions in block *i* before all its pair frequencies fall to one, and let *r*_*i*_ be its residual length. Then *h*_*i*_ +*r*_*i*_ −1 = *u* (*n*_*i*_). A one-symbol block also satisfies this identity because *n*_*i*_ = 1, *h*_*i*_ = 0, and *r*_*i*_ = 1. After all blocks reach this stage, the entire sequence has length 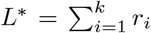 and lies in the frequency-one regime. Lemma 1 therefore gives exactly *L*^*^ − 1 further substitutions. Adding both stages yields

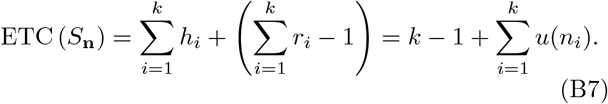

#### b. The grouped-run sequence exceeding the bound

Now we show that *S*_**n**_ exceeds the bound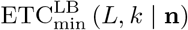.

Write

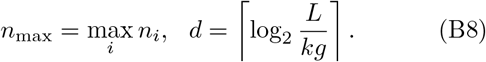

The definition of *u* gives *u*(*x*) ≥ ⌈log_2_ *x*⌉ for every positive integer *x*. Also, *n*_max_ ≥ *L/k* ≥ *L/kg*. Therefore,

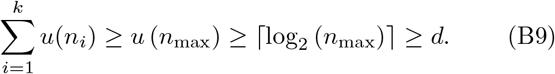

If at least two symbol counts exceed one, the first inequality is strict and ETC (*S*_**n**_) *>* ETC^LB^ (*L, k* | **n**) because the largest block contributes *u*(*n*_max_) and another block contributes at least one more.

It remains to consider compositions of the form **n** = (*r*^*^, 1, …, 1), with the largest count in any labeled position. Here *g* = 1, *L* = *r*^*^ + *k* − 1, and *n*_max_ = *r*^*^ ≥ 3. We only need to show *u* (*r*^*^) *> d*.

If *r*^*^ is not a power of two, put *q* = ⌊log_2_ *r*^*^⌋. Its binary expansion has at least two ones, so *u* (*r*^*^) is at least *q* + 1. Therefore,

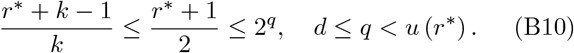

If *r*^*^ = 2^*q*^, then *q* ≥ 2 and *u* (*r*^*^) = *q*. For *k* ≥ 3,

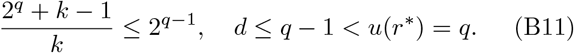

Thus, ETC (*S*_**n**_) is strictly above the bound in these cases as well.

#### c. A separate ordering for the binary exception

The only cases left are *k* = 2 with **n** = (2^*q*^, 1) or **n** = (1, 2^*q*^ ). The grouped-run sequence equals the lower bound in these cases. For **n** = (2^*q*^, 1), we choose a different ordering:

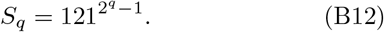

The trailing run has length 2^*q*^ − 1 and contributes *u* (2^*q*^ − 1) = 2*q* − 2. The initial 12 adds two symbols to the final frequency-one residual. Consequently,

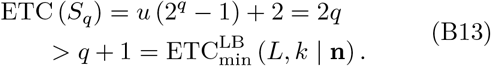

For the reversed composition **n** = (1, 2^*q*^ ), use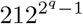.

The same run-length calculation gives ETC = 2*q*.

Every admissible **n**_*S*_ therefore contains a sequence strictly above its lower bound.

#### d. Consequences for the reference means

Taking the uniform average gives

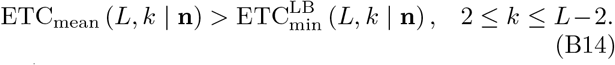

As shown in Section II.B, the global ensemble mean is the sequence-count-weighted average of the composition-conditioned ensemble means. Because the weights are positive and sum to one,

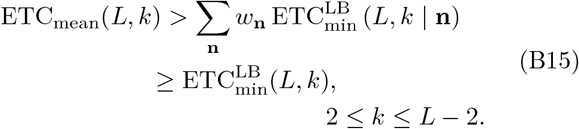

The remaining cases are degenerate. For *k* = *L* and *k* = *L* − 1, every sequence has ETC *L* − 1, and lower bounds also equal *L* − 1. The denominators are zero there. For *k* = 1, every sequence is constant and ETC is zero.

### 4. Relation to the true minimum

For fixed *k*, the grouped-run sequence 1^*L*−*k*+1^2 3 · · · *k* and the identity in Appendix B.3 give

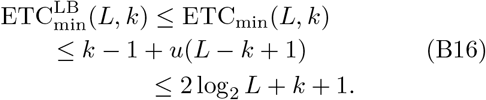

Thus, the certified global lower bound and the true global minimum are both *O*_*k*_ (log *L*).

To compare the resulting global coordinates, let true minimum *TM* = ETC_min_(*L, k*), certified lower bound 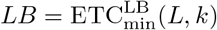, and mean ETC *µ* = ETC_mean_(*L, k*).

Define

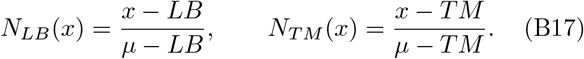

#### Proposition 4

**(Global lower-anchor error)** *For each fixed integer k* ≥ 2, *both denominators in Eq*. (B17) *are positive for sufficiently large L. For such L, the two global coordinates satisfy*

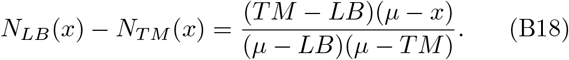

*Their difference has the uniform bound*

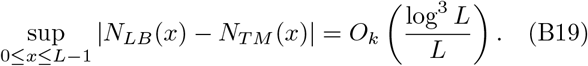

Theorem 1 gives *µ* = Θ_*k*_ (*L/* log *L*). Equation (B16) gives *LB, TM* = *O*_*k*_ (log *L*). Hence both denominators are positive and of order *L/* log *L* for sufficiently large *L*. Direct subtraction gives Eq. (B18). Since 0 ≤ *µ* ≤ *L* − 1, we have |*µ* − *x*| ≤ *L* − 1 throughout 0 ≤ *x* ≤ *L* − 1. Together with *TM* − *LB* = *O*_*k*_(log *L*), these bounds give Eq. (B19).

Using the certified lower reference instead of the true minimum therefore changes the global coordinate by a quantity that tends to zero at fixed *k*.

## Appendix C: Exact ceiling of maximum ETC

To reach ETC *L* − 1, every substitution must shorten the sequence by exactly one position. If the first selected pair occurs twice or more, that step already removes too many positions. Conversely, if *m*(*S*) equals one, Lemma 1 gives exactly *L* − 1 substitutions. Thus, attaining the ceiling is equivalent to having no initial pair with more than one nonoverlapping occurrence.

For a given *k*, there are *k*(*k* − 1) unequal-symbol pairs. Equal-symbol pairs require a separate count because their occurrences can overlap. For example, *aaa* contains two adjacent aa positions but only one nonoverlapping occurrence. Write *r*_*j*_ for the lengths of the separate maximal runs of symbol *a*, where a maximal run is a consecutive block that cannot be extended by another *a*. Then

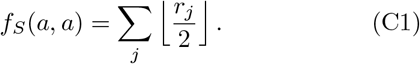

If this sum is at most one, at most one run can have length two or three, and every other run has length one. The pair *aa* can therefore occupy at most two adjacent positions. Summing over the *k* equal-symbol pair types gives at most 2*k* positions. Together with the unequal pairs, this yields

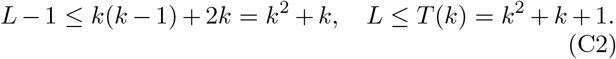

For attainment, start from a de Bruijn sequence of order two: a cyclic sequence in which each of the *k*^2^ ordered pairs appears exactly once. Appending its first symbol gives a linear sequence of length *k*^2^ + 1 with the same pair property. Replace each unique *aa* occurrence by *aaa*. This adds one position per symbol without repeating any unequal pair. The resulting sequence has length *T* (*k*) = *k*^2^ + *k* + 1, and every pair still has nonoverlapping frequency one.

To cover *k* ≤ *L* ≤ *T* (*k*), choose the de Bruijn sequence to start with 1, 2, …, *k*. This prefix can be prescribed because removing the path 1 → 2 → · · · → *k* from the directed de Bruijn graph leaves an Euler trail from *k* to 1. Then extend every *aa* to *aaa*. The constructed sequence also has ETC = *T* (*k*) − 1. Any prefix with *k* ≤ *L* ≤ *T* (*k*) still contains all *k* symbols. Taking a prefix cannot increase a pair frequency, so each such prefix also attains its own ceiling. This proves

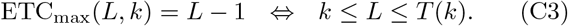

## Appendix D: The scaling of maximum ETC and ensemble mean

The lower bound comes from counting how many sequences can be represented by a small number of substitutions. The upper bound applies to every sequence and limits how many substitutions can remain after repeated pairs become infrequent.

### 1. Asymptotic lower bound for global-ensemble mean ETC

We count sequences with small ETC by counting the records needed to reconstruct them. Let *E* be a nonnegative integer, and let *N*_*E*_(*L, k*) be the number of length-*L* sequences over a fixed *k*-alphabet whose ETC is at most *E*. A sequence with ETC at most *E* can be reconstructed using at most *E* substitution rules. We will use this count to show that sequences with sufficiently small ETC form a vanishing fraction of the global ensemble as *L* increases. For this upper count, sequences may omit alphabet symbols. Including them can only increase the count relative to Ω_*L,k*_.

To see what a reconstruction rule means, suppose one substitution replaces the ordered pair *xy* by a fresh symbol *z*. We store the reverse instruction *z* → *xy*. A collection of these instructions is called a binary grammar [25]. Here, it is simply a record of the substitutions already performed by NSRPS [5].

For example, the deterministic substitutions for the sequence 001001 are

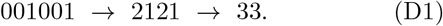

The corresponding reconstruction rules are

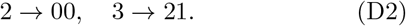

Expanding one terminal 3 gives 21 and then the original-symbol block *P* = 001. The terminal sequence has two copies of 3, so the full input is *S* = *P* ^2^. Thus the substitution record and the terminal multiplicity recover the original sequence exactly.

In general, write the terminal constant sequence as *a*^*v*^, where *a* is the terminal symbol. Expanding through the recorded rules gives a block *P*, and *S* = *P*^*v*^. The number of rules needed to expand *a* equals ETC(*S*) = *e*. Because the total length *L* is fixed, the multiplicity is forced to be *v* = *L/*|*P* | . We therefore need to count only the records that reconstruct *P* from *a*, without a separate choice for *v*.

We represent each reconstruction record as a partial parse tree, retaining only the first expansion of each rule [26]. Begin at *a*. Whenever a rule is expanded, draw two ordered branches leading to its left and right symbols. An original symbol ends a branch. If the same rule is encountered again, retain its symbol as a leaf without expanding its descendants. This construction preserves the information needed for reconstruction while avoiding repeated copies of the same subtree.

For our count, traverse the tree depth first, visiting the left branch before the right branch. Assign each rule a reference number when it is first encountered. These reference numbers are independent of the original symbol labels. Each leaf then records either an original symbol or a reference to a previously encountered rule. Fig. D1 illustrates how a repeated subtree is replaced by a reference leaf.

We now count these trees in two parts. First count their shapes, including the left-to-right order of the branches. The number of shapes with *e* internal nodes and *e* + 1 leaves is the Catalan number *C*_*e*_ = (2*e*)!*/*[(*e* + 1)!*e*!]. Next choose what each leaf records. There are *k* original symbols and at most *e* rule references, giving at most *k* +*e* choices per leaf and at most (*k* + *e*)^*e*+1^ assignments for a given shape. Allowing all these choices includes invalid references, but that only increases the upper count.

Every valid record determines one block *P*. Records for which |P| does not divide *L* can be discarded. Different records may reconstruct the same sequence, so counting records can overcount sequences but cannot miss any sequence with ETC at most *E*. Summing over 0 ≤ *e* ≤ *E* gives

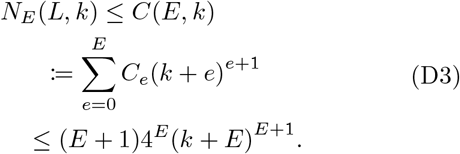

The final inequality uses *C*_*e*_ ≤ 4^*e*^ and bounds each of the *E* +1 summands by the same largest expression. Thus *N*_*E*_(*L, k*) is bounded by a quantity determined only by *E* and *k*. The notation *C*(*E, k*) denotes this upper count of reconstruction records. We next compare it with the number of sequences in the global ensemble.

For any 0 *< ε <* 1, choose the cutoff

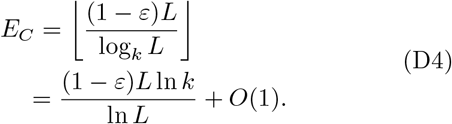

Substituting this cutoff gives

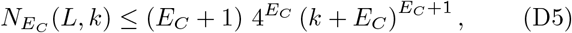

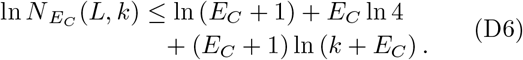

At fixed *k*, the largest term in the right-hand side is *E*_*C*_ln *E*_*C*_ = (1 − *ε*)*L* ln *k* + *o*(*L*). Therefore

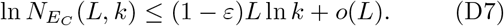

At fixed *k*, at most *k*(*k* − 1)^*L*^ length-*L* sequences have missing symbols. Their ratio is

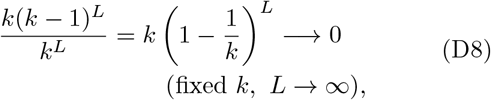

and |Ω_*L,k*_| = *k*^*L*^ (1 − *o*(1)) and ln |Ω_*L,k*_| = *L* ln *k* +*o*(*L*). The count of sequences below the cutoff *E*_*C*_ therefore grows exponentially more slowly, so

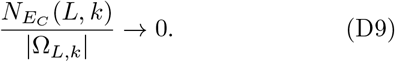

A uniformly sampled global-ensemble sequence consequently has ETC greater than *E*_*C*_ with probability tending to one. Since ETC is integer-valued,

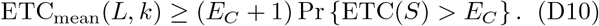

Taking *L* to infinity and then *ε* ↓ 0 yields

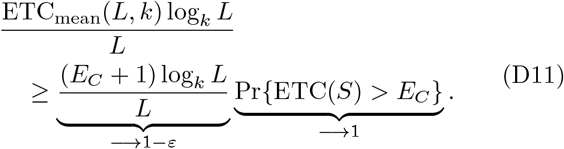

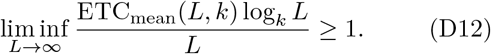

### 2. Asymptotic upper bound for maximum ETC

For the upper bound, we associate each current symbol with the sequence of original symbols that it represents.

#### a. Unique recovery of current-symbol blocks

At any substitution step, if two current-symbol blocks have the same original expansion, they are identical. Assume that it holds immediately before a substitution of *xy* by a fresh symbol *z*. Take two blocks after that substitution and expand each *z* back to *xy*. The resulting blocks have the same original expansion and are therefore identical by the assumption. To complete the induction, we must show that each such preceding block has only one representation after substitution.

**FIG. D1.**
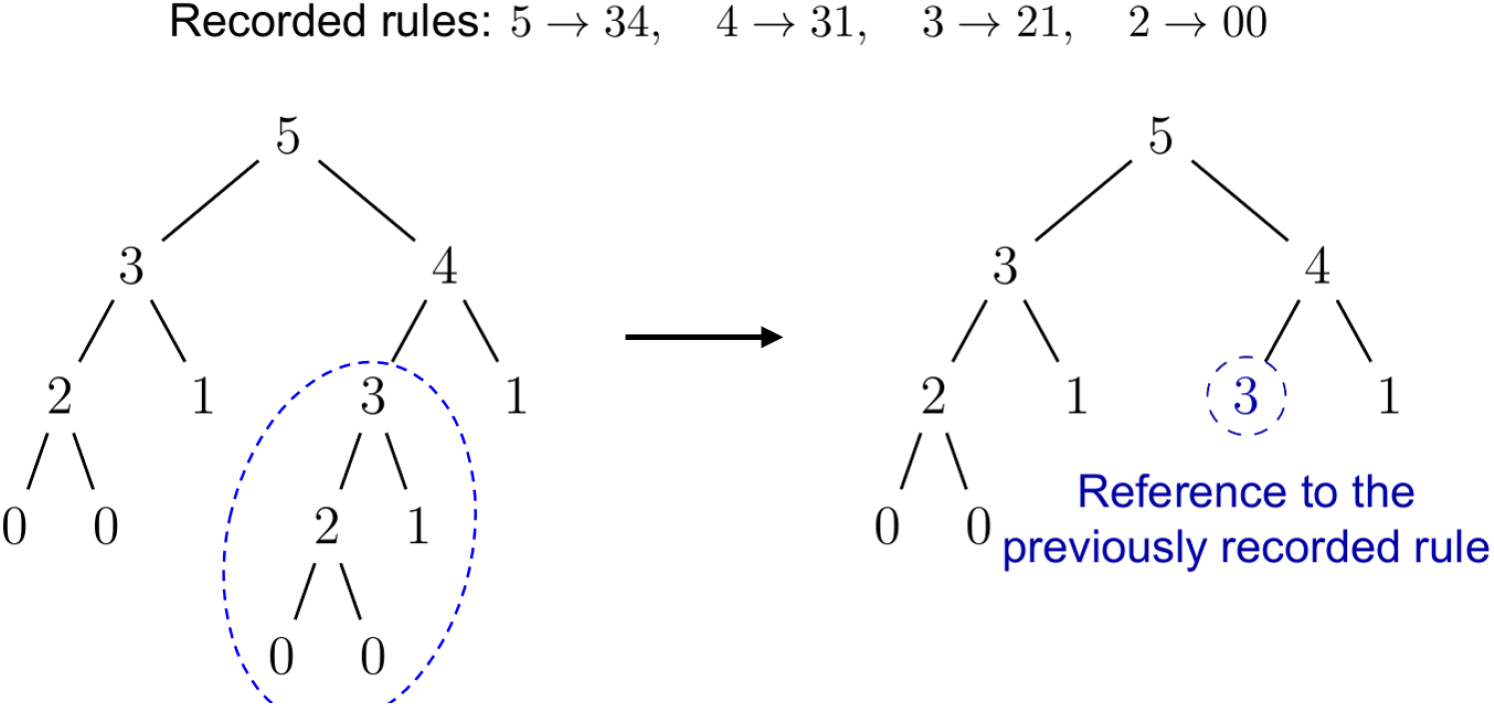
Partial parse tree representation of substitution rules. Left: Full expansion of the recorded rules reconstructs the block *S* = 0010011. Right: Each rule is expanded only at its first occurrence. The repeated subtree rooted at symbol 3 (dashed ellipse) is replaced by a leaf referencing the previously recorded rule (dashed circle). The resulting tree has *e* = 4 internal nodes and *e* + 1 = 5 leaves and retains all information needed to reconstruct *S*.

If *x* and *y* are different, occurrences of *xy* cannot overlap. Their locations uniquely determine where *z* appears because the substitution replaces every occurrence. If *x* and *y* are equal, each maximal run transforms as

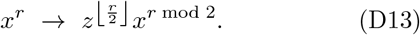

A contiguous block within the transformed run consists of copies of *z* followed by at most one *x*. Its expansion length fixes both the number of *z* symbols and whether the final *x* is present. A block crossing several runs has a unique decomposition because other symbols separate the runs. This statement applies to two-symbol blocks, i.e., pairs.

#### b. Bounding the number of remaining substitutions

Choose an integer cutoff *B* ≥ 1. Follow the substitutions until the sequence is constant or maximum pair frequency becomes *m*(*S*) ≤ *B*. Every earlier substitution decreases the length by at least *B* + 1, so there are at most ⌊(*L* − 1)*/*(*B* + 1)⌋ pre-cutoff steps.

If the residual sequence is nonconstant, denote its current length by *L*^*^. Split it into ⌊*L*^*^*/*2⌋ disjoint adjacent pairs, leaving the last symbol unpaired if needed. Expand each pair into its original-symbol sequence. These expanded phrases occupy disjoint parts of the original sequence, and their total length is at most *L*.

By the recovery uniqueness proved above, copies of the same expanded phrase must come from the same ordered current pair. There can be at most *B* copies of such pair in the disjoint grouping because *m*(*S*) is at most *B*.

We choose another integer cutoff *q* ≥ 2 for the length of expanded phrase. An expanded phrase is *short* if it contains at most *q* original symbols and *long* otherwise. There are at most 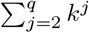 possible *short phrases* and their total contribution is at most *B* times that count. Every *long* phrase uses at least *q* + 1 original positions, so there are at most ⌊*L/*(*q* + 1)⌋ *long phrases*. Hence

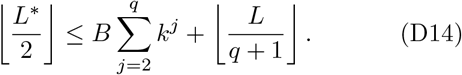

At most *L*^*^ − 1 ≤ 2⌊*L*^*^*/*2⌋ post-cutoff steps remain. Adding the pre-cutoff steps, and retaining the conventional ceiling, gives the sequence-independent bound

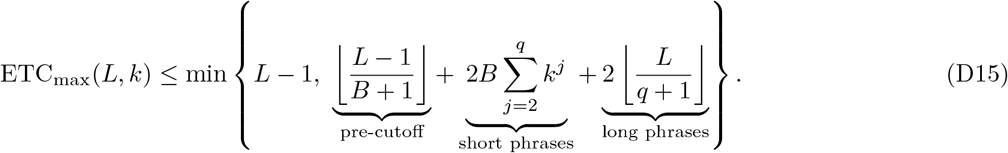

For the fixed-*k* asymptotic bound, set *λ* = log_*k*_ *L* and choose

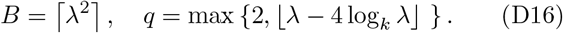

For sufficiently large *L* at fixed *k, λ* − 4 log_*k*_ *λ* ≥ 2, so *q* = ⌊*λ* − 4 log_*k*_ *λ*⌋.

The pre-cutoff term is *O*_*k*_ (*L/λ*^2^ ). The *short-phrase* term has the same order because 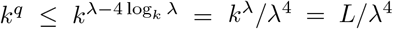 and *B* has order *λ*^2^. Since *q/λ* → 1, the *long-phrase* term is (2 + *o*(1)) *L/λ*. Since *L* − 1 is generally unattainable (Appendix C), combining this upper bound with the preceding ensemble-mean lower bound gives

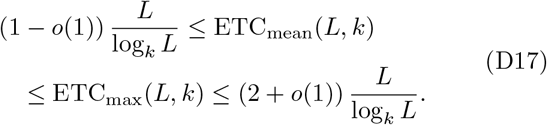

### 3. Composition-conditioned mean scaling

For a positive probability vector **p** = (*p*_1_, …, *p*_*k*_) with 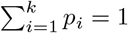, define the normalized entropy as in Section III.C:

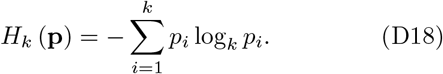

#### Proposition 5

**(Composition-conditioned means)**

*Fix an integer k* ≥ 2 *and a probability vector* **p** *with p*_*i*_ *>* 0 *for every symbol. For each integer L* ≥ *k, let* **n**^(*L*)^ *be a positive integer composition with*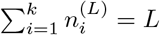. *If /L* → **p**, *then*

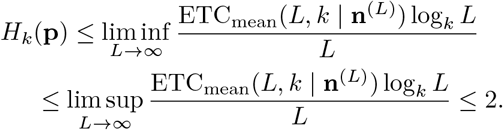

To prove the lower bound, we apply the counting argument of Appendix D.1 to this composition class. Stirling’s formula gives

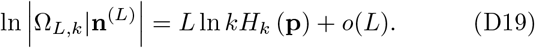

Therefore, the logarithm of the composition class size has leading term *L* ln *kH*_*k*_ (**p**), rather than the global-ensemble term *L* ln *k*. The factor *H*_*k*_ (**p**) accounts for the size of the composition class.

To obtain a lower bound for the mean within this composition class, fix 0 *< ε <* 1 and choose the integer cutoff

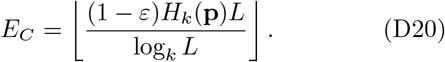

The previously derived quantity 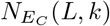 counts all length-*L* sequences with ETC at most *E*_*C*_, regardless of composition. It therefore also bounds the number of such sequences in this particular composition class.

Substituting the chosen *E*_*C*_ into that bound gives

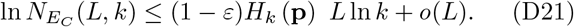

For a sequence *S* uniformly sampled from the composition class, dividing this count by the corresponding class size bounds the probability of ETC(*S*) ≤ *E*_*C*_ as

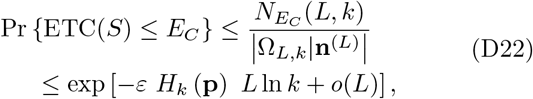

whose limit is zero. Thus, Pr {ETC(*S*) *> E*_*C*_} tends to one. The mean therefore satisfies

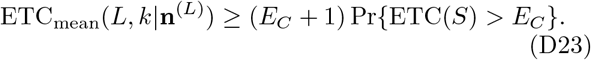

Multiply both sides by log_*k*_ *L/L*. The probability factor tends to one, while the cutoff factor satisfies

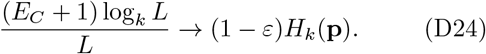

Taking *L* to infinity and then *ε* ↓ 0 yields the lower bound

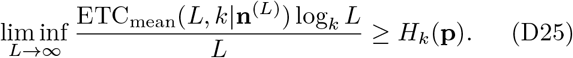

The upper bound in Appendix D.2 holds for every sequence and therefore also bounds each composition-conditioned mean. This gives the upper bound in Proposition 5. Since *H*_*k*_(**p**) *>* 0, the mean has order *L/* log_*k*_ *L* for fixed *k* and **p**.

## Appendix E: Greedy construction

### 1. Extension algorithm

The constructive search begins from an exact threshold witness at *L* = *T* (*k*). The extension code resumes from the longest sequence in the saved history. Let *S* be the current sequence of length *L* and let *e* = ETC(*S*). Each extension cycle starts with insertion block length *n* = 1 and seeks a candidate with ETC at least *e* + 1.

For a one-symbol insertion, the candidate set includes each of the *k* symbols at every position 0, …, *L*. Position 0 is before the first symbol and position *L* is after the last. The ranking score is the largest adjacent-pair count in the resulting sequence. These counts include overlapping occurrences, unlike the nonoverlapping frequencies used by ETC. The code scores all candidates without computing their ETC.

Candidates are evaluated in increasing score order. Ties are resolved first by numerical symbol order and then by insertion position from left to right. The code computes ETC for one candidate at a time and accepts the first with ETC at least *e* + 1. It does not evaluate the remaining candidates after a success. If ETC values tie, it keeps the first candidate in the evaluation order.

If no one-symbol candidate reaches the target, the search tries block lengths 2 through 11 in increasing order. For each length *n*, it selects the min(*n*+1, *k*) least frequent symbols in *S*, resolving equal symbol counts by numerical label. It forms every length-*n* block from this set, allowing repeated symbols, and scores each block at every insertion position. Candidates are ranked by increasing score, then by lexicographic block order, then by insertion position. Lexicographic order compares numerical labels at the first position where two blocks differ. ETC evaluation again stops at the first candidate that reaches the target.

The current sequence *S* and target *e* + 1 remain fixed throughout these searches. If every candidate through insertion block length 11 fails to reach the target, the code selects the retained one-symbol candidate with the highest ETC. This fallback increases length by one even though ETC can stay unchanged or decrease. The block-length cutoff limits the computational cost.

After either a successful insertion or the fallback, the selected sequence becomes *S*. The code recomputes its ETC before saving the sequence. The next extension cycle uses the updated *e* and restarts with *n* = 1. Full flowchart is shown in Fig. E1.

## Appendix F: Potts-chain generation and calibration

### 1. Stationary chain and entropy rate

Let *x* and *y* index the symbol labels and let *f* denote the favored symbol. The coupling *J* controls the preference for equal neighboring symbols, while the field *η* favors *f* . We use a transfer matrix representation of the one-dimensional Potts chain [15]. For *J* ≥ 0, the symmetric transfer matrix is

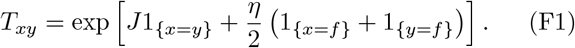

Each indicator equals one when its stated condition holds and zero otherwise. The field contribution is split equally between the two ends of each adjacent pair.

Let *λ* be its largest eigenvalue and *v* a positive eigenvector. Define

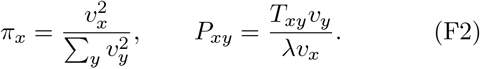

Then *P* is a stochastic matrix, *πP* = *π*, and *π*_*x*_*P*_*xy*_ = *π*_*y*_*P*_*yx*_. Sequences start with *X*_1_ ∼ *π* and continue by sampling *X*_*t*+1_ ∼ *P*_*X*_ . This directly samples a stationary chain without a burn-in period.

Using natural logarithms, define

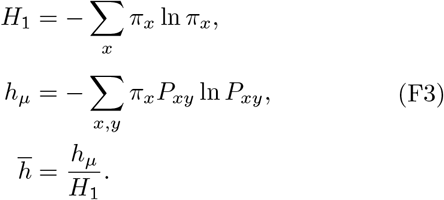

At *J* = 0, the transition rows equal *π* and the process is iid for every field. Therefore, 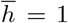 even when the marginal composition is nonuniform. The joint probabilities shown in Fig. 5 are *π*_*x*_*P*_*xy*_.

### 2. Joint length and alphabet size calibration

In a zero-field Potts chain, *π*_*x*_ = 1*/k* and

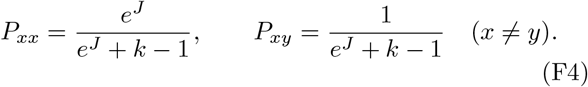

Because *H*_1_ = ln *k*, the normalized entropy rate reduces to

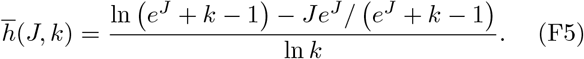

For numerical calibration, we reparametrized *J* using

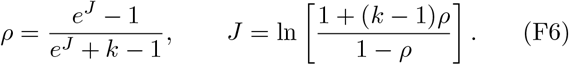

For each *k* and target 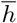, we solved 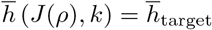 for *ρ* ∈ [0, 1). Fig. 4 used the full factorial grid *L* ∈ {2000, 4000, 8000, 16000}, *k* ∈ {2, 5, 10, 20, 50}, and 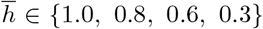. Each cell contained 200 independent process sequences. Sequences were rejected and redrawn if they omitted any alphabet symbol.

**FIG. E1.**
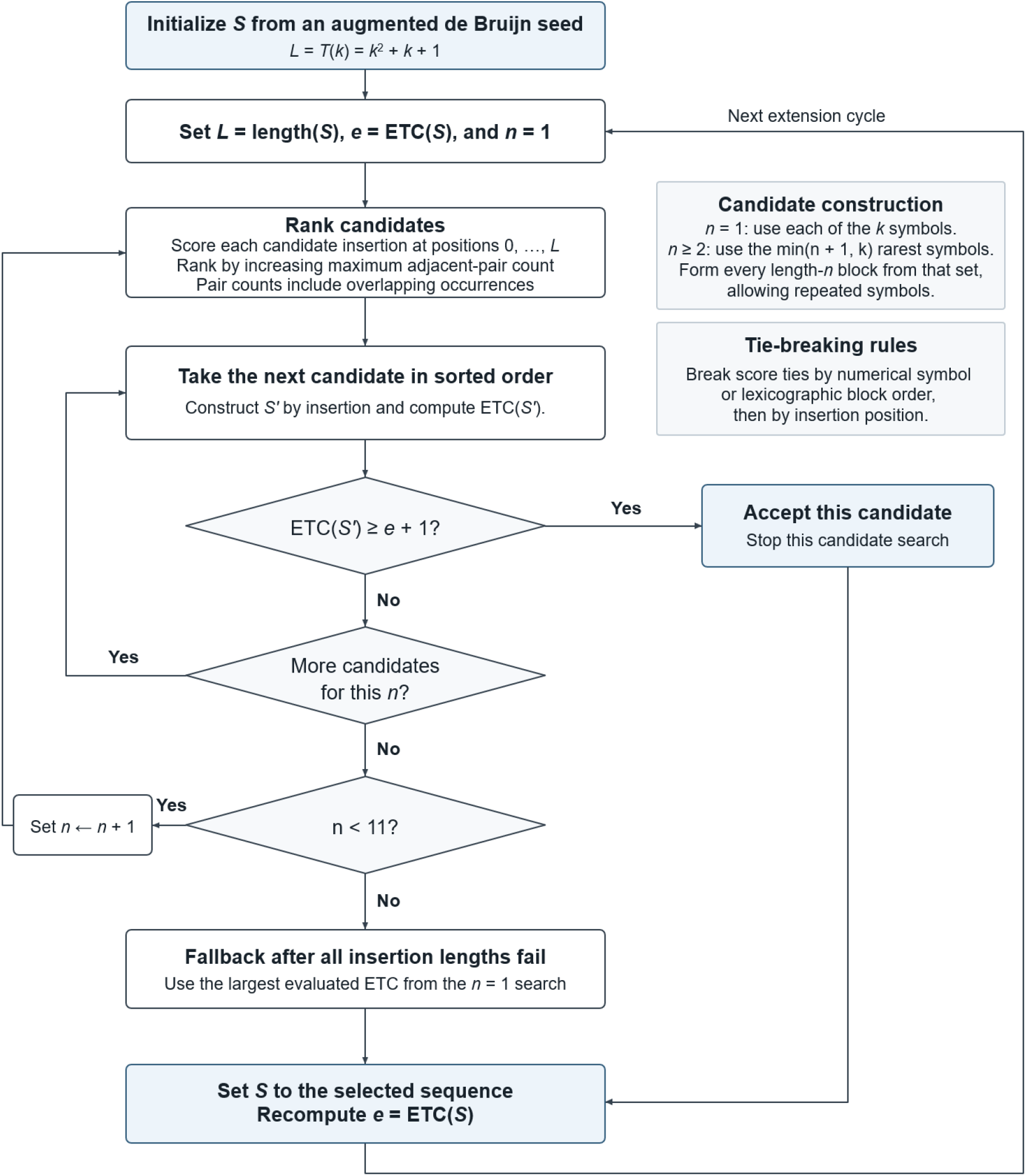
Flowchart of algorithm for greedy construction of high-ETC sequences.

### 3. Composition and entropy-rate matching

Fig. 5 fixed *L* = 2000 and *k* = 5. The target favored-symbol probabilities were *p*_*f*_ ∈ {0.2, 0.35, 0.5, 0.65, 0.8} and the target normalized entropy rates were 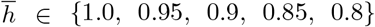. For each of the 25 cells, *J* and *η* are chosen jointly to solve

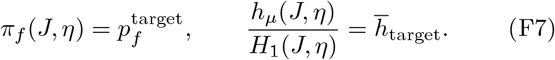

At 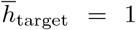, the solution is *J* = 0 and *η* = ln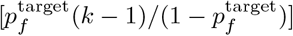. Otherwise, bounded non-linear least squares searches 0 ≤ *J, η* ≤ 20 from fixed multiple starting points and retains the smallest residual. Each cell contained 200 independent stationary sequences.

## Appendix G: Genomic sequence analysis

### 1. Sequence sources and selection

We used the UCSC hg38 chromosome 22 reference sequence [27]. CpG island annotations came from cpg-IslandExt [28], and LINE annotations came from RepeatMasker [29]. The frozen UCSC cpgIslandExt, RepeatMasker, and simpleRepeat annotation tables had an October 18, 2022 snapshot date. Coding regions came from GENCODE release 50 on GRCh38.p14 [30]. We retained coding DNA sequence (CDS) entries from protein-coding MANE Select transcripts [31], so the exon category excludes untranslated regions. The simpleRepeat track contains annotations generated by Tandem Repeats Finder [32] and provided the tandem repeat exclusion mask.

CpG island intervals were retained as annotated. Overlapping RepeatMasker calls with repeat class LINE were merged. CpG and LINE sequences used the reference-genome orientation. Coding exons were individual CDS intervals rather than concatenated transcripts. CDS sequences on the negative strand were reverse complemented. The CDS phase was not used to trim codon boundaries because each nucleotide, not each codon, was analyzed as a symbol. Sequences shorter than 200 bases or containing non-ACGT characters were excluded. Sequences that overlapped with a tandem repeat or either of the other two target annotation classes were excluded. For each retained interval, we selected one nonoverlapping background interval of exactly the same length outside all target annotations, tandem repeats, and non-ACGT runs. Background allocation proceeded from longer to shorter intervals and searched the source 5-Mb block before expanding to neighboring blocks. Previously selected backgrounds were excluded from subsequent draws. The backgrounds used the same orientation as the corresponding matching sequences.

The final sample contained 285 CpG islands, 7,938 LINE intervals, 428 coding exons, and 8,651 unique matched backgrounds. The retained lengths (mean ± SD) were 626 ± 442 bp for CpG islands (range: 201–2,552 bp), 531 ± 575 bp for LINEs (200–7,533 bp), and 402 ± 472 bp for coding exons (200–5,655 bp).

### 2. Complexity features and reference sampling

We encoded A, C, G, and T as 0, 1, 2, and 3 and computed ETC. The global ensemble reference used *k* = 4 at each observed length. Its mean was estimated from 100 independent uniform four-symbol sequences, rejecting draws that omitted a symbol. The composition reference preserved the ordered nucleotide-count vector (*n*_A_, *n*_C_, *n*_G_, *n*_T_). For each distinct vector, 100 uniform permutations of the corresponding ACGT multiset estimated the reference mean. Counts were neither sorted nor reassigned to different labels.

We also computed the LZ76 phrase count *c*_76_ and applied the conventional length normalization with alphabet size four [33, 34]:

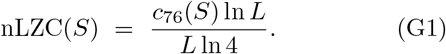

### 3. Length-adjusted annotation comparisons

For each annotation and each complexity feature, we fitted an ordinary least-squares model to the annotation sequences and their own matched backgrounds:

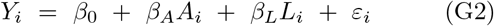

where *Y*_*i*_ is the feature value, *A*_*i*_ equals one for the annotation and zero for background, and *L*_*i*_ is sequence length. Length entered linearly without an interaction. Annotation effects were evaluated with two-sided *t* tests using conventional standard errors under an independent-error assumption. Benjamini–Hochberg FDR adjustment was applied jointly to the 12 annotation-effect tests (three annotations × four complexity features; Table G1).

### 4. Binary classification and evaluation

We evaluated three binary tasks, each comparing one annotation with its own matched background. Each of the four features entered a separate ridge logistic regression as a single predictor. Features were standardized using training data only. Five outer folds provided one out-of-fold probability per sequence. Within each outer training set, three-fold grouped cross-validation selected *C* from 0.01, 0.1, 1, 10, and 100 by AUROC. Groups were defined from the annotation sequences’ 5-Mb chromosome blocks. Each matched background was assigned to its annotation partner’s group, and exons from the same gene were kept together. Five background intervals lay outside their partner’s block. The folds were therefore not strictly geographically separated. We used a maximum of 5,000 iterations. ROC curves and AUROCs pooled the out-of-fold probabilities across the five outer folds.

**TABLE G1.** Length-adjusted annotation effects and FDR-adjusted *p*-values.

| Annotation | Complexity feature | $\beta_A$ | $t$ (df) | $p_{\text{FDR}}$ |
| --- | --- | --- | --- | --- |
| CpG island | nLZC | -0.025235 | -8.329 (567) | < 0.0001 |
|  | nETC | -0.020665 | -7.902 (567) | < 0.0001 |
|  | cETC | 0.024586 | 11.611 (567) | < 0.0001 |
|  | nETC - cETC | -0.045251 | -26.006 (567) | < 0.0001 |
| LINE | nLZC | -0.012245 | -23.505 (15,873) | < 0.0001 |
|  | nETC | -0.009488 | -21.180 (15,873) | < 0.0001 |
|  | cETC | 0.015467 | 37.033 (15,873) | < 0.0001 |
|  | nETC - cETC | -0.024955 | -82.763 (15,873) | < 0.0001 |
| Coding exon | nLZC | 0.000539 | 0.210 (853) | 0.8339 |
|  | nETC | -0.007671 | -3.327 (853) | 0.0011 |
|  | cETC | -0.002429 | -1.220 (853) | 0.2430 |
|  | nETC - cETC | -0.005243 | -4.866 (853) | < 0.0001 |

## Notes

### Competing Interest Statement

The authors have declared no competing interest.

